# Identification of SLC25A46 interaction interfaces with mitochondrial membrane fusogens Opa1 and Mfn2

**DOI:** 10.1101/2023.12.29.573615

**Authors:** Sivakumar Boopathy, Bridget E. Luce, Camila Makhlouta Lugo, Pusparanee Hakim, Julie McDonald, Ha Lin Kim, Jackeline Ponce, Beatrix M. Ueberheide, Luke H. Chao

**Affiliations:** Department of Molecular Biology, Massachusetts General Hospital, Boston MA 02114, USA; Department of Genetics, Harvard Medical School, Boston MA 02115, USA; Proteomics Resource Center, Division of Advanced Research Technologies, New York University Langone Health Center, New York NY 10016, USA; Department of Biochemistry and Molecular Pharmacology, New York University Langone Health Center, New York NY 10016, USA

**Keywords:** Mitochondria, Membrane fusion, GTPase, Structural model, Protein cross-linking, Mass spectrometry, Protein-protein interaction, Mitochondrial solute carrier

## Abstract

Mitochondrial fusion requires the sequential merger of four bilayers to two. The outer-membrane solute carrier protein SLC25A46 interacts with both the outer and inner-membrane dynamin family GTPases Mfn1/2 and Opa1. While SLC25A46 levels are known to affect mitochondrial morphology, how SLC25A46 interacts with Mfn1/2 and Opa1 to regulate membrane fusion is not understood. In this study, we use crosslinking mass-spectrometry and AlphaFold 2 modeling to identify interfaces mediating a SLC25A46 interactions with Opa1 and Mfn2. We reveal that the bundle signaling element of Opa1 interacts with SLC25A46, and present evidence of a Mfn2 interaction involving the SLC25A46 cytosolic face. We validate these newly identified interaction interfaces and show that they play a role in mitochondrial network maintenance.

## Introduction

Mitochondrial fission and fusion are membrane remodeling processes required for maintenance of a healthy mitochondrial reticulum (1, 2), which regulates the number and distribution of mitochondrial DNA and respiratory chain proteins (1-5). A distinct feature of mitochondrial fusion is the sequential fusion of two pairs of bilayers. Key questions regarding the spatiotemporal control of these events remain. Specifically, how are the activities of the outer and inner membrane fusion GTPases, Mfn1/2 and Opa1 respectively, coordinated and regulated?

SLC25A46 interacts with both fusion GTPases and has emerged as an inter-membrane bridging protein, playing roles in regulating mitochondrial membrane dynamics (6-9). SLC25A46 is a member of the solute carrier family 25 of mitochondrial transporters, which transport substrates such as nucleotides, vitamins, amino acids, fatty acids across the inner mitochondrial membrane (IMM) (10). Residing in the outer mitochondrial membrane (OMM), SLC25A46 lacks the signature sequence motifs of the SLC25 family (10), does not have an identified substrate and contains a distinct N-terminal extension. Mutations in SLC25A46 result in optic atrophy spectrum disorder, Leigh syndrome and pontocerebellar hypoplasia (11-25).

In SLC25A46 knockout and knockdown models of human neurological disease, mitochondria are often enlarged and aggregated in nervous tissues, exhibiting detached, vesiculated cristae (22, 25-29). Respiratory chain complex activity is impaired in these models (26, 27), and ATP levels are reduced (27, 28). In human cell lines, knockdown of SLC25A46 induces a tubular mitochondrial network whereas overexpression results in mitochondrial fragmentation (6, 9, 25), suggesting a role in downregulating fusion. The phenotypes of SLC25A46 knockout varied depending on the cell lines tested, displaying mitochondrial fragmentation in human fibroblasts and HeLa cells (8), and hyper-filamentous network in other cell lines (30, 31). Knockout of the yeast orthologue Ugo1 (∼22% sequence identity) also induced fragmentation (32-34). These observations argue for a role of SLC25A46 in regulating mitochondrial fusion where SLC25A46 levels at the OMM are critical. In fibroblasts and iPSC-derived cerebral neurons, SLC25A46 is focally present at tips and branches of mitochondrial tubules and at sites of fusion and fission (8). Interestingly the SLC25A46 puncta were proximal to Opa1 in human fibroblasts (8), suggesting spatially-defined interactions with SLC25A46 play a role in fusion regulation.

Understanding the molecular details of the SLC25A46-Mfn2-Opa1 interactions can provide constraints on how SLC25A46 organizes sites for mitochondrial membrane remodeling and inform mechanisms for coordinated OMM/IMM fusion. In yeast, Ugo1 physically interacts with Fzo1 and Mgm1, the orthologous fusion GTPases (34-37). Previous studies indicate Opa1 and Mfn2 interacts with SLC25A46 (6-9), however the specific interfaces mediating these interactions was undefined. In this study, we used crosslinking-mass spectrometry and AlphaFold 2 modeling to identify the protein-protein interaction surfaces of SLC25A46-Opa1 and SLC25A46-Mfn2. We validate this newly identified SLC25A46-Opa1 interface by mutagenesis and demonstrate a role for this interface in mitochondrial network maintenance.

## Results

### SLC25A46 interacts with Opa1 via its bundle signaling element

Previous studies used crosslinking, immunoprecipitation and mass spectrometry to identify an interaction network of Opa1, Mfn1/2 and SLC25A46 (6-9). We over-expressed human SLC25A46 with Opa1 in *Pichia pastoris* under the control of a strong promoter (alcohol oxidase) (**Figure 1A**) and found that detergent (GDN)-solubilized SLC25A46 in *P. pastoris* lysate interacts with column-immobilized Opa1 from lysate in the absence of crosslinkers (**Figure 1B**). Likewise, column-immobilized SLC25A46 was also found to interact with Opa1 (**Supporting Figure 1A**).

**Figure 1.**
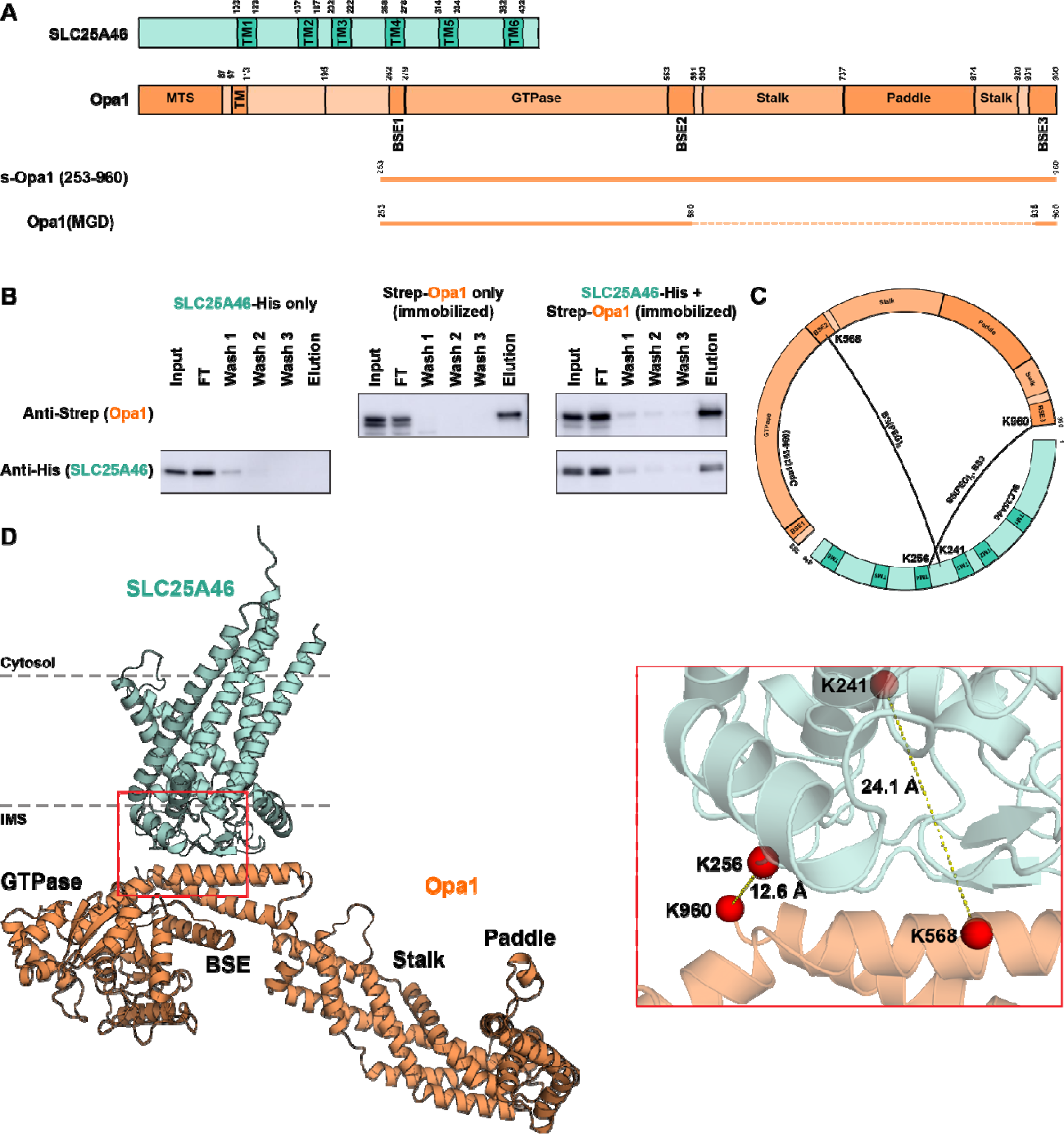
SLC25A46 interacts with the bundle signaling element (BSE) of Opa1. A. Domain diagrams of SLC25A46 and Opa1. s-Opa1 (amino acids 253-960) and Opa1(MGD) (minimal GTPase domain comprising the GTPase domain and three BSE helices) are shown as line diagrams below full length Opa1 sequence. B. GDN extracts prepared from *P. pastoris* expressing SLC25A46-His, Strep-Opa1 or co-expressing SLC25A46-His and Strep-Opa1 subjected to StrepTactin column binding and elution. C. Crosslinks identified in BSE helices 2 and 3 of Opa1. Circular map of SLC25A46 and Opa1 with identified lysine-lysine crosslinks (black arcs). Crosslinker used to generate the crosslink is indicated. D. AlphaFold 2 model of the SLC25A46-Opa1 interaction. N-terminal region of SLC25A46 comprising residues 1-83 (disordered) not displayed, residues 1-263 of Opa1 not modeled. Box region displaying identified crosslinks and the calculated C - C distances based on the model.

To identify the specific interface responsible for the SLC25A46-Opa1 interaction, we crosslinked the purified complex for analysis by mass spectrometry. Using bis(sulfosuccinimidyl)suberate (BS3) we identified a crosslink between K241 of SLC25A46 and K568 of Opa1 bundle signaling element (BSE) helix 2. Intralinks consistent with folded Opa1 secondary structure elements were identified (**Supporting Figure 1B**). A second crosslinker Bis-N-succinimidyl-(pentaethylene glycol) (BS(PEG)5) with a larger spacer length also crosslinked K241(SLC25A46)-K568(Opa1), and identified a second crosslink between K256 of SLC25A46 and the K960 of Opa1 BSE helix 3 (**Figure 1C**). We modeled this interaction using Alphafold 2 (38, 39), which consistently revealed a high-confidence and robust interface mediated by the Opa1 BSE, with a topology consistent with the proteins’ sub-organellar localization (6, 22, 25, 40, 41) (**Figure 1D, Supporting Figure 1C**).

### Validation of SLC25A46:Opa1-BSE interaction

Examination of the SLC25A46-Opa1 interface in the AlphaFold 2 model revealed two salt bridges comprising residues R257 (SLC25A46) and E561 (Opa1); R347 (SLC25A46) and D565 (Opa1). Adjacent to the salt bridges, a cluster of hydrophobic residues, I229, I230, I349, Y357, V359 and L360 in SLC25A46 and A569, F572 and T576 in Opa1 BSE helix 2 appear to further stabilize the interaction (**Figure 2A**). We tested this newly identified SLC25A46-Opa1 interface by mutating the salt bridges and hydrophobic site residues in SLC25A46 and evaluating if the interaction with Opa1 was maintained. While recombinantly expressed and purified WT SLC25A46 bound column-immobilized Opa1 (consistent with observations in lysates), both the I349D mutation and the R257A/R347A double mutation diminished the interaction (**Figure 2B and Supporting Figure 2A**).

**Figure 2.**
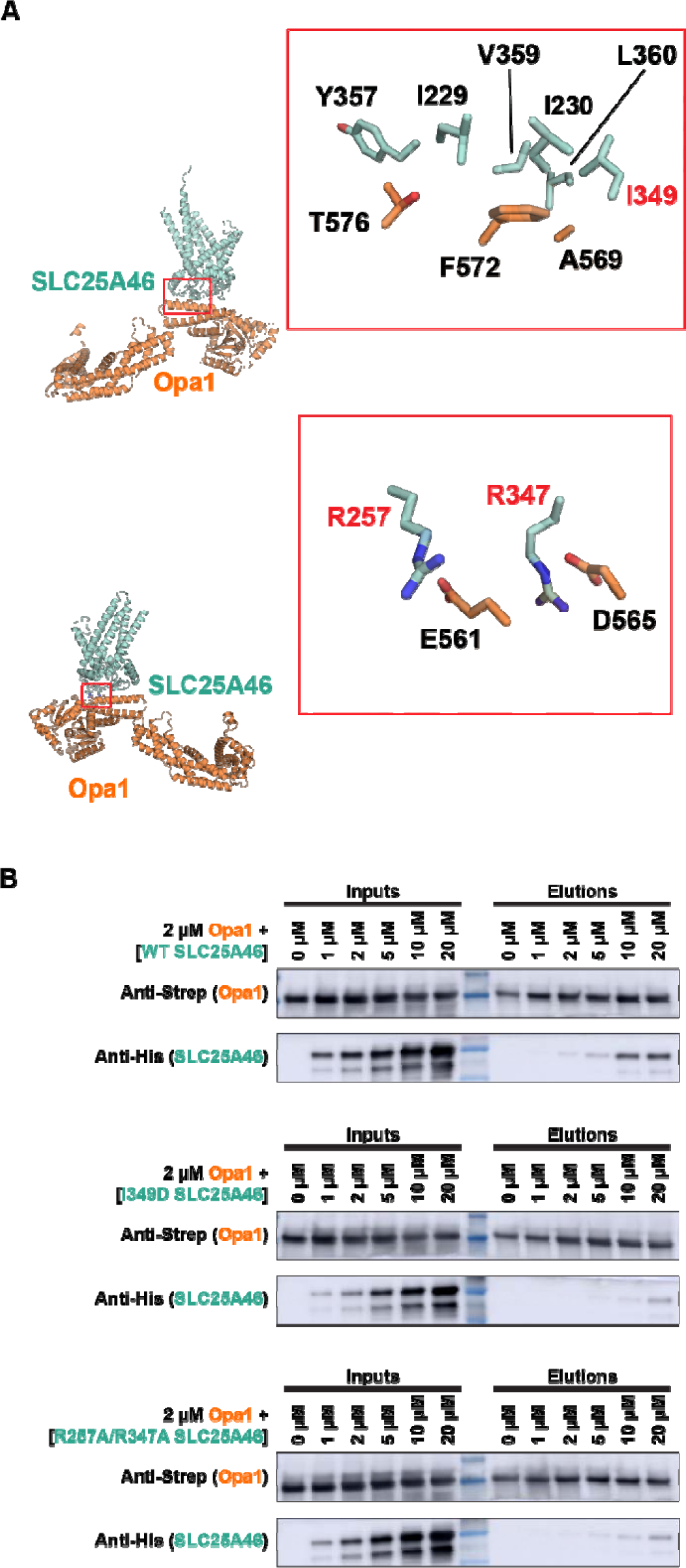
SLC25A46 mutations in the binding interface diminish binding to Opa1. A. SLC25A46-Opa1 interface observed in AlphaFold 2 model. Top view: Adjacent to the salt bridges is a cluster of hydrophobic residues comprising of I229, I230, I349, Y357, V359, L360 of SLC25A46 and T576, F572, A569 of Opa1. Bottom view: Predicted salt-bridge interactions between R257 of SLC25A46 and E561 of Opa1, and R347 of SLC25A46 and D565 of Opa1. Interface residues highlighted. Mutated residues indicated in red. B. Purified SLC25A46-His WT, I349D mutant or R257A/R347A double mutant were incubated with 2 µM of Strep-Opa1 and subjected to StrepTactin binding and elution experiments. Input and elution analyzed by Western blotting.

The BSE is essential for transducing GTP-hydrolysis stimulated activities of dynamin related protein (42, 43). In dynamin, the BSE frequently moves closer to the GTPase upon GTP hydrolysis (44-46). To determine if the nucleotide-state of Opa1 affects its interaction with SLC25A46, we incubated the lysates with GTPγS, GMPPCP and GDP prior to affinity capture on StrepTactin resin. We found that the presence of nucleotide or nucleotide analogues did not affect co-elution, indicating nucleotide-coupled BSE rearrangement does not affect Opa1-SLC25A46 interaction (**Supporting Figure 2B**).

We also generated a minimal construct consisting of the GTPase domain and the BSE alone (Opa1(MGD)), to test that the “paddle” and stalk self-assembly regions are dispensable for the Opa1-SLC25A46 interaction (**Figure 1A**). SLC25A46 co-elutes with Opa1(MGD), indicating an Opa1(MGD) construct is sufficient for binding (**Supporting Figure 3A**). We found the same two crosslinks observed for Opa1-SLC25A46 between Opa1(MGD) and SLC25A46. In addition, we observed a third crosslink between K387 of SLC25A46 and K568 of Opa1 in the same vicinity, strengthening our findings (**Supporting Figure 3B and 3C**).

**Figure 3.**
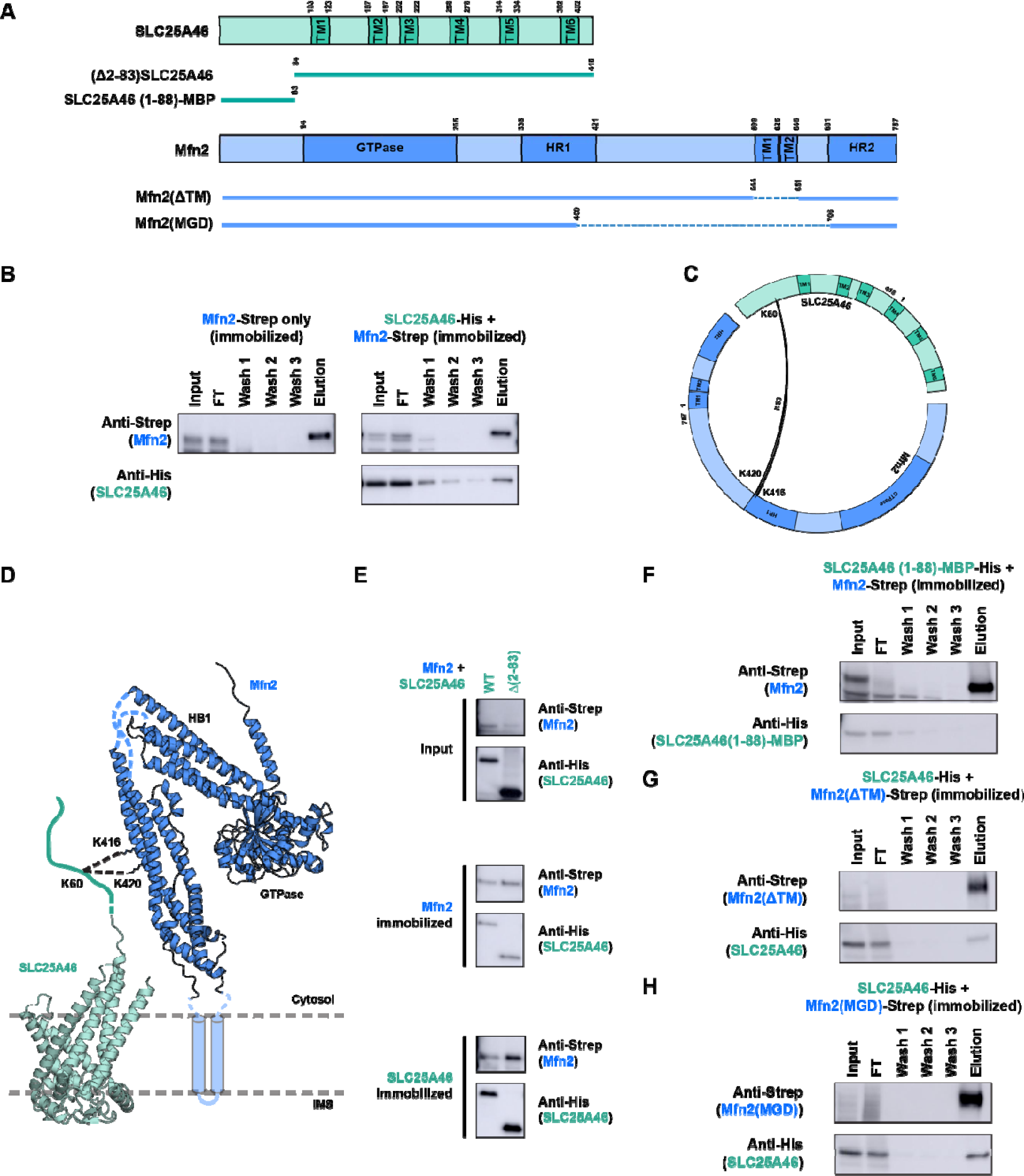
SLC25A46 interactions with Mfn2. A. Domain diagrams of full length and truncated constructs of SLC25A46 and Mfn2 are depicted. B. SLC25A46-His co-elutes upon elution of bound Mfn2-Strep. GDN extracts prepared from *P. pastoris* expressing Mfn2-Strep or co-expressing SLC25A46-His and Mfn2-Strep were subjected to StrepTactin column binding and elution. C. Circular map of SLC25A46 and Mfn2 with the identified lysine-lysine crosslinks (black arcs). Crosslinker used is indicated. Crosslinks were identified between the cytosol-exposed N-terminal K60 of SLC25A46 and the HR1 lysines K416 and K420 of Mfn2. D. Model of the SLC25A46-Mfn2 interaction. The identified crosslinks are depicted as dotted lines in the AlphaFold 2 generated models of SLC25A46 and Mfn2. Helical bundle 1 (HB1) indicated. The N-terminal region of SLC25A46 comprising residues 1-83 is predicted to be disordered and is displayed as a hand-drawn cartoon. The transmembrane helices of Mfn2 were poorly modeled and are shown as transparent cylinders, to indicate uncertainty. E. (Δ2-83)SLC25A46-His co-elutes upon elution of bound Mfn2-Strep. GDN extracts prepared from *P. pastoris* expressing Mfn2-Strep and co-expressing SLC25A46-His or (Δ2-83)SLC25A46-His were subjected to StrepTactin column binding and elution. F. Residues 1-88 of SLC25A46 fused to MBP and His tag at the C-terminus (SLC25A46(1-88)-MBP-His) does not co-elute with immobilized Mfn2-Strep. G. A Mfn2 lacking the two transmembrane segments (Mfn2(ΔTM)-Strep) co-elutes with SLC25A46-His. H. A minimal Mfn2 construct comprising helical bundle 1 (HB1) and the GTPase domain co-elutes with SLC25A46-His.

### Cytosolic facing SLC25A46 interactions with the Mfn2 ectodomain

We next investigated if SLC25A46 also interacts with the outer membrane fusion protein Mfn2. We co-expressed the two proteins in *P. pastoris* (**Figure 3A**) and subjected the GDN-solubilized lysates to column-binding experiments. SLC25A46 co-elutes upon elution of column-immobilized full-length Mfn2 (**Figure 3B**), and Mfn2 co-elutes with resin bound SLC25A46 (**Supporting Figure 4A**). To test the SLC25A46-Mfn2 interaction was nucleotide-state dependent, we treated lysates with GTPγS, GMPPCP or GDP during detergent extraction. The presence of these nucleotides did not enhance or diminish interaction of Mfn2 with SLC25A46 (**Supporting Figure 4B**).

**Figure 4.**
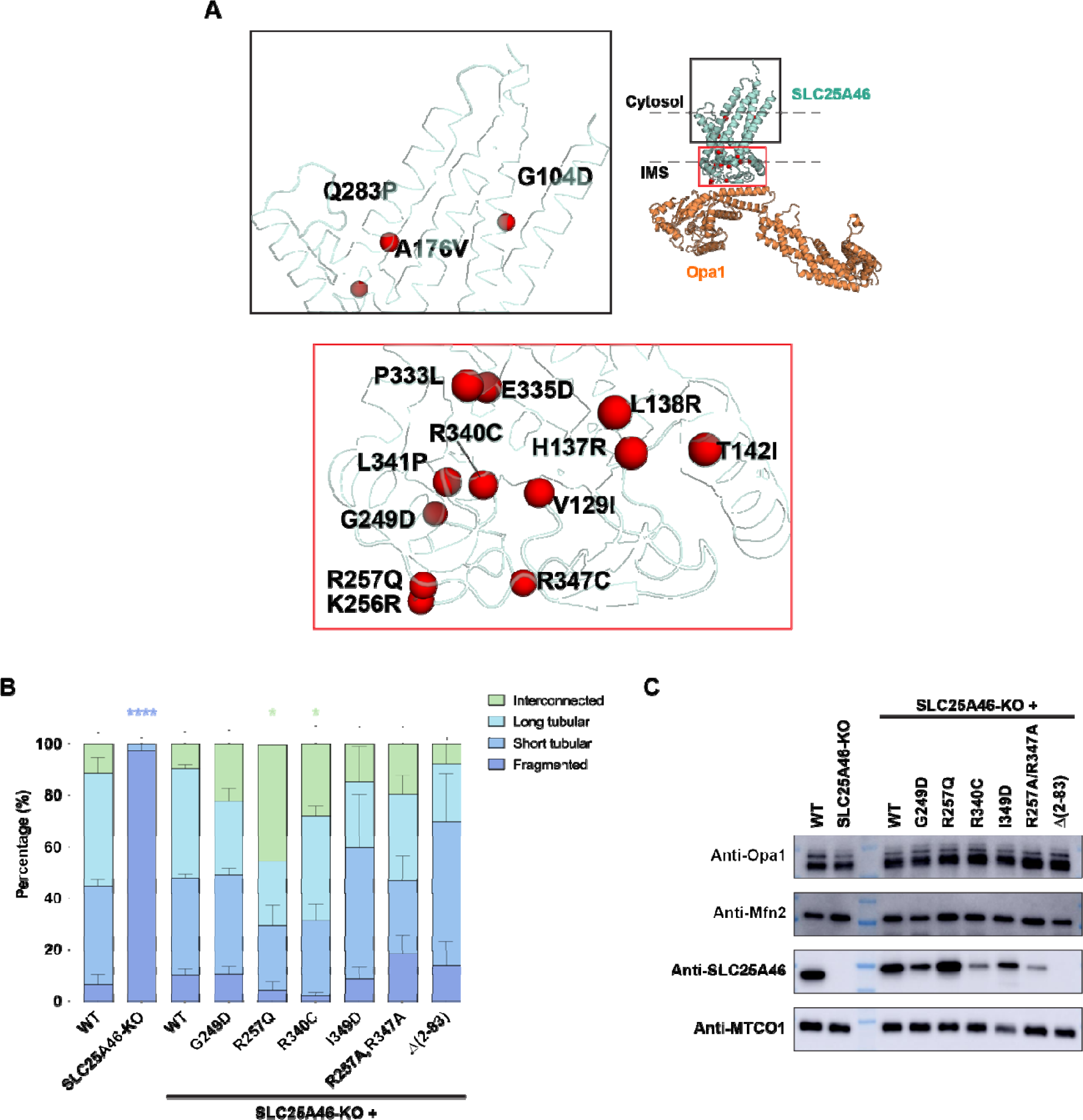
Mutations disrupting Opa1 interaction alter mitochondrial morphology. A. Identified SLC25A46 disease mutations mapped on AlphaFold 2 model of SLC25A46-Opa1 complex. Box regions enlarged on the right to display the mutations. B. Graph bar representing the relative proportion of scored mitochondrial morphologies in HCT116 WT, SLC25A46-KO or SLC25A46-KO cells expressing SLC25A46 variants. Mitochondrial morphology for 300 cells were scored for each condition and evaluated (see Materials and Methods). Significance of difference is tested relative to wild type using unpaired t test with Welch’s correction. Blue ****: p<0.0001 when compared to scored WT cells showing fragmented mitochondria. Green *: p<0.05 when compared to scored WT cells showing interconnected mitochondrial network. C. Western blot of expression levels for Opa1, Mfn2, SLC25A46 or MTCO1 in HCT116 WT, SLC25A46-KO or SLC25A46-KO cells expressing SLC25A46 variants. (Δ2-83)SLC25A46 not detected as commercially available antibodies detect the N-terminal epitope of SLC25A46. MTCO1 is used as loading control.

To gain insight into the nature of the SLC25A46-Mfn2 interaction, we performed crosslinking and mass-spectrometry on the purified complex. In addition to intramolecular crosslinks consistent with folded Mfn2 (**Supporting Figure 4C**), two inter-links were identified – between K60 residue of SLC25A46 and residues K416 and K420 of Mfn2 (**Figure 3C**). K60 of SLC25A46 is part of the first ∼83 residues that are predicted to be disordered in the cytosol (**Figure 3D**). Residues K416 and K420 are part of the cytosol facing heptad repeat 1 (HR1) of Mfn2 (**Figure 3D**). In contrast to the Opa1-SLC25A46 models, AlphaFold 2 predicted interface was low confidence for the SLC25A46-Mfn2 complex, particularly for the N-terminal region of SLC25A46, and the Mfn2 transmembrane segments (**Supporting Figure 4D**).

Since crosslinks were observed at K60 in SLC25A46, we tested the role of these N-terminal residues in binding Mfn2. We deleted the first 82 amino acids of SLC25A46 termed (Δ2-83)SLC25A46, and co-expressed this protein with Mfn2 in *P. pastoris*. Interestingly, (Δ2-83)SLC25A46 still co-eluted with Mfn2 in the affinity capture experiments (**Figure 3E**), suggesting interactions outside this N-terminal segment mediating Mfn2 interactions. Consistent with this idea, MBP-fusion of the SLC25A46 N-terminus (SLC25A46(1-88)-MBP) was insufficient to capture Mfn2 (**Figure 3F**). To exclude the possibility that the transmembrane segments of Mfn2 interact with SLC25A46, we co-expressed SLC25A46 with Mfn2 lacking the transmembrane helices (Mfn2(ΔTM)). We observed co-elution of SLC25A46 with Mfn2(ΔTM) (**Figure 3G**), supporting an interaction between the Mfn2 ectodomain and cytosol-exposed regions of SLC25A46. Expression of a minimal Mfn2(MGD) construct, containing only the helical bundle (HB1) and GTPase domain (47), still bound SLC25A46, suggesting this region of the ectodomain interacts with the SLC25A46 cytosolic face (**Figure 3H**).

### Mutations disrupting SLC25A46-Opa1 interaction alter mitochondrial morphology

A cluster of disease mutations reported in SLC25A46 face the IMS (**Figure 4A**) (16). While many mutations were reported to destabilize the protein, certain disease-causing mutations (T142I and R340C) cause a hyperfused mitochondrial phenotype (6, 8, 9, 16, 25), similar to that observed with siRNA/shRNA knockdown of SLC25A46 (6, 9, 25).

To investigate cellular mitochondrial phenotypes upon SLC25A46 perturbations, we imaged mitochondrial networks in the human colorectal carcinoma cell line HCT116. Under steady-state conditions, WT and SLC25A46 KO HCT116 cells demonstrate similar mitochondrial network morphology. Previous studies show that nutrient starvation triggered stress can induce mitochondrial hyperfusion (48, 49). To test if mitochondria in HCT116 cells undergo similar stress-induced change, cells were grown in serum-free medium. Under such conditions, we observed significant mitochondrial fragmentation in SLC25A46 KO cells but not in WT cells. Reintroduction of WT SLC25A46 rescued this phenotype (**Figure 4B, Supporting Video 1**), consistent with previous observations in human fibroblasts and HeLa cells (8). We found that expression of R257Q or R340C SLC25A46 mutants induced interconnected mitochondrial networks, also in agreement with previous studies (8, 25) (**Figure 4B, Supporting Video 1**). The mitochondrial morphology in G249D-expressing cells was similar to WT (**Figure 4B, Supporting Video 1**). Expression levels of R340C and G249D mutants never reached WT-like levels (**Figure 4C**), likely due to protein instability (6, 8, 9, 16, 22).

We next tested the effects of the newly identified Opa1-SLC25A46 interactions on mitochondrial morphology. Expression of I349D SLC25A46 (disrupts a hydrophobic interface between the Opa1 BSE) produced shorter mitochondrial tubules compared to WT, whereas R257A/R347A mutant (disrupt SLC25A46-Opa1 salt bridge interactions) resulted in more fragmented mitochondria (**Figure 4B, Supporting Video 1**). These results suggest that the identified SLC24A45-Opa1 interaction interface is important for maintaining a tubular mitochondrial morphology. Expression of the (Δ2-83)SLC25A46 resulted in an increase in short mitochondrial tubules (**Figure 4B, Supporting Video 1**). In these experiments, we did not observe changes in levels of Mfn2 or Opa1 due to SLC25A46 knockout or expression of varying levels of mutant SLC25A46 (**Figure 4C**). The expression level of (Δ2-83)SLC25A46 could not be evaluated due to the inability of commercial antibodies to detect this mutant (**Figure 4C**).

We also investigated SLC25A46 disease mutants expressed in yeast. Under these conditions, many of SLC25A46 mutants still bound Opa1 (**Supporting Figure 5B**). We also saw little effect of patient-derived mutations on the Mfn2-SLC25A46 interaction (**Supporting Figure 5B**). We note that in these binding experiments we were unable to achieve similar expression levels of the respective proteins, and binding may only describe one aspect a complex phenotype only observable under cellular conditions.

## Discussion

Mitochondria undergo morphological changes tightly coupled to cellular physiology (4). While the identity of the GTPases responsible for sequential mitochondrial fusion have long been identified (50), and mechanisms for individual fusion steps have been described (51-54), questions remain regarding how outer and inner-membrane activities are coordinated and regulated. Emerging evidence points to SLC25A46 as a regulator of fusion activity, with protein levels influencing the fusion-fission balance (6, 8, 25). Consistent with this role, SLC25A46 is observed to localize to the fusion sites (8). In this study, we investigate the physical interaction between SLC25A46 and the membrane fusogens Opa1 and Mfn1/2. Combining crosslinking mass-spectrometry, and AlphaFold 2 modeling, we identify and validate the BSE region in Opa1 interaction with the IMS-face of SLC25A46. We also present evidence for an interaction between SLC25A46 cytosolic face and the Mfn2 ectodomain. The Mfn2 region we identify includes helical bundle 1 (HB1) and GTPase domains, but not the transmembrane segments. This is consistent with yeast studies, where residues 1-294 of Ugo1 interacts with Fzo1 in a region comprising residues N-terminal to the first transmembrane helix and the C-terminal helical repeat 2 (34).

Reports from several studies indicate that mitochondrial morphology is sensitive to SLC25A46 levels. Schuettpelz *et al.* reported mitochondrial fragmentation in human fibroblasts and HeLa cells upon SLC25A46 KO (8, 55). In SLC25A46 KO HCT116 cells, we observed mitochondrial fragmentation upon serum starvation. In addition, we observed a ∼2-fold slower growth of this cell line, consistent with observations in SLC25A46 deleted fibroblasts (8). In previous studies, overexpression of SLC25A46 increased mitochondrial fragmentation, suggesting that at high levels, SLC25A46 may downregulate the activities of the protein fusogens. (6, 25, 31). One possible mode of inhibitory action may be SLC25A46 binding resulting in steric competition for Opa1 self-assembly, which relies on the BSE and stalk regions to mediate formation of helical filaments (56-58). Consistent with this prediction, levels of Opa1 oligomeric complexes were found to increase upon loss of SLC25A46 (8).

In our study, SLC25A46 mutants targeting Opa1 interaction did not promote a hyperfused phenotype, but instead disrupted the mitochondrial network. This suggests that a SLC25A46-Opa1 interaction promotes fusion necessary for a tubular network. Although we could not estimate the relative levels of (Δ2-83)SLC25A46, this N-terminal deletion reduced the filamentous network, while increasing the proportion of short tubules, also suggesting a fusion defect. Since the SLC25A46 N-terminal region alone is insufficient to bind with Mfn2, we speculate that this region may play a stabilizing role facilitating interactions of additional cytosolic-facing regions with the ectodomain of Mfn2. Taken together, these observations provide evidence for SLC25A46 interactions playing an activating role in fusion when expressed at physiological levels.

Our results provide a basis for future functional analyses of SLC25A46 regulation of the fusion machinery. SLC25A46 protein-protein interactions also may regulate cristae morphology, which is dramatically affected upon loss of SLC25A46 (6, 8, 25-27, 29). Proteomics studies have also implicated SLC25A46 endoplasmic reticulum interaction partners in maintaining ER-mitochondria contacts facilitating lipid flux (6-8). Future mechanistic structure-activity studies are necessary to explore these other roles of SLC25A46 in cristae regulation and lipid metabolism.

## Experimental procedures

### Plasmids

Genes for SLC25A46, Mfn2, Opa1(253-960), Opa1(MGD) constructs were synthesized (GenScript) and cloned into pPICZ A for *P. pastoris* expression. C-terminus of SLC25A46 was fused to a (G_4_S)_3_ linker followed by TEV cleavage site and deca-histidine. N-terminus of Opa1(253-960), Opa1(263-960), Opa1(MGD) was tagged with Twin-Strep tag followed by HRV 3C cleavage site and (G_4_S)_3_ linker. The C-terminus of Mfn2 was fused to G_3_S linker, HRV 3C cleavage site and Twin-Strep tag. For co-expression, SLC25A46 and either Mfn2 or Opa1 with tags with its own AOX1 promoter and terminator were cloned into the same plasmid. (SLC25A46(1-88)-MBP) was generated from SLC25A46(1-88) fused to MBP followed by a GGGS linker TEV site and His10 tag. Mfn2(ΔTM) was generated from Mfn2(2-544)-GGS linker-Mfn2(651-757). Mfn2(MGD) was generated from Mfn2(2-400:706-757). Strep-Opa1(253-960) and Strep-Opa1(MGD) were also cloned into pET-28a(+) vector for expression in *E. coli*.

### Generation of *Pichia* strains

Expression strains were generated from *Pichia pastoris* strain SMD1163 (*his4 pep4 prb1*) (kind gift from Dr. Tom Rapaport (Harvard Medical School) according to established protocols (59). Plasmids were transformed into SMD1163 by electroporation and grown on YPDS (1% yeast extract, 2% peptone, 2% dextrose, 1 M sorbitol) zeocin (InvivoGen, ant-zn-1p) plates. 7-14 resulting colonies were grown in BMGY (1% yeast extract, 2% peptone, 100 mM potassium phosphate pH 6.0, 1.34% yeast nitrogen base, 4 × 10^−5^% biotin, 1% glycerol) and induced in BMMY (replaced glycerol with 0.5% methanol in BMGY). Cultures were screened for protein expression by Western using anti-His, anti-Strep, anti-SLC25A46, anti-Opa1 or anti-Mfn2 antibodies.

### Binding experiments using lysates

*P. pastoris* (1 L) expressing SLC25A46, Opa1, Mfn2 alone or in combination were harvested and cryo-milled using SPEX SamplePrep 6875D Freezer/Mill. For StrepTactin column-binding and elution, milled cells (125 mg) were resuspended in 3 ml of 25 mM HEPES, 150 mM NaCl, pH 8.0 buffer containing 1% GDN and protease inhibitors, tumbled for 1-1:30 h at 4 °C. Milled cell input was increased to 375 mg (3x original input) when needed for increased signal. Suspension was centrifuged at 20,000 for 30 min at 4 °C. Supernatant (100 µl) was incubated with 50 µl of StrepTactin XT 4Flow high-capacity resin in a spin column (Abcam, ab286861) at 4°C for 15 min followed by centrifugation at 500 xg for 30 s. Resin was washed 3x with 100 µl buffer containing 0.1% GDN. To elute, the resin was incubated with 100 µl buffer with 50 mM biotin for 15 min and centrifuged. Same procedure for Ni-NTA binding experiments except that the binding and elution buffer contained 80 mM and 500 mM imidazole, respectively.

### AlphaFold 2 modeling

Protein sequences for human SLC25A46, Opa1 and Mfn2 were run through in a AlphaFold2 Google ColabFold – v 1.5.3 workbook (38, 39, 60). Five independent runs generated the top three ranked relaxed models for SLC25A46-Opa1 and SLC25A46-Mfn2 models respectively.

## Data availability

Mass spectrometry data available on request.

## Supporting information

This article contains supporting information.

## Acknowledgements

We thank Dr. Yuan Gao and Dr. Tom Rapoport for providing *Pichia pastoris* strain SMD1163. We thank all the members of the Chao lab for critical feedback.

## Funding and additional information

Mass spectrometric experiments supported by a shared instrumentation grant from the NIH, 1S10OD010582-01A1 towards purchase of Orbitrap Eclipse. This work was supported by funding from the National Institutes of Health (R35GM142553 to L.H.C.).

## Abbreviations and nomenclature

OMM: outer mitochondrial membrane
IMM: inner mitochondrial membrane
SLC: solute carrier family
SLC25A46: solute carrier family 25 member A46
Opa1: optic atrophy 1
Mfn1/2: mitofusin 1/2
iPSC: induced pluoripotent stem cells
BSE: bundle signaling element
HR1/2: heptad repeat 1/2
ER: endoplasmic reticulum
MGD: minimal GTPase domain
DDM: n-Dodecyl-β-D-Maltopyranoside
GDN: glyco-diosgenin
PA: phosphatidic acid
TM: transmembrane segment
MTS: mitochondrial targeting sequence
IMS: inter-membrane space

