## Supporting Data for "Identification of SLC25A46 interaction interfaces with mitochondrial membrane fusogens Opa1 and Mfn2"

**Supporting Information**

**Supporting experimental procedures**

**Binding experiments using lysates – additional details**

For nucleotide treatments, StrepTactin binding experiments were performed as described but the buffer used was 25 mM HEPES, 100 mM NaCl, 50 mM KCl, 10 mM MgCl_2_.6H_2_O, pH 8.0, 1% GDN (extraction) or 0.1% GDN (binding and elution) and containing 2 mM of GTPγS (Sigma-Aldrich, 10220647001), GMPPCP (Sigma-Aldrich, M3509-25MG) or GDP (Sigma-Aldrich, G7127-25MG).

To assess binding of SLC25A46-His disease mutants to Opa1 or Mfn2, *P. pastoris* strains co-expressing the proteins were cultured in 5-10 ml BMMY. Equal number of cells (estimated by OD600) were harvested from each culture. The cells were resuspended in 0.5 ml of 25 mM HEPES, 150 mM NaCl, pH 8.0 containing protease inhibitors. An equal volume of 0.5 mm glass beads was added and lysed using a Retsch TissueLyzer. To the recovered lysate, GDN was added from a stock solution of 10% to a final concentration of 1% and incubated in the cold room for 1.5 h tumbling gently. The lysate was then centrifuged at 21,100 xg for 30 min at 4 °C. 150-300 ul of the lysate was applied to either Ni-NTA (Bio-Rad, 780-0800) or StrepTactin XT 4Flow resin pre-equilibrated with the above buffer containing 10 mM imidazole (for Ni-NTA) or 1 mM EDTA (StrepTactin) and 0.1% GDN. Resins were washed 3x and eluted with buffer containing 500 mM imidazole (for Ni-NTA) or 50 mM biotin (StrepTactin) and 0.1% GDN. The samples were subjected to western blotting using anti-Strep (Mfn2 or Opa1) and anti-His (SLC25A46) antibodies.

**Protein expression**

For protein expression in *P. pastoris*, a single desired clone was inoculated in 5-10 ml BMGY and then scaled up to 500 ml BMGY media and grown overnight at 30 °C. The culture was harvested and resuspended in 6 litres BMMY at an OD600 of 1.0 for protein induction. The induced cultures were then grown at 20 °C and harvested 24 h later. Cell pellets were frozen in liquid nitrogen and stored at –80 °C.

Opa1 and Opa1(MGD) were also purified from *E. coli*. For expression in *E. coli*, transformed BL21-CodonPlus (DE3)-RIL cells (Agilent, 230245) were grown in LB at 37 °C to an OD600 of 0.7 and induced with 0.2 mM IPTG (Gold Biotechnology, I2481C200) and grown overnight at 16 °C. Cultures were harvested and stored at –80 °C until use.

**Purification of SLC25A46**

Harvested *P. pastoris* cells expressing WT, I349D mutant or R257A/R347A double mutant were milled under cryogenic conditions using SPEX SamplePrep 6875D Freezer/Mill. All purification steps were done at 4 °C. The milled cells were resuspended in 50 mM HEPES, 250 mM NaCl, 50 mM imidazole, pH 7.8 containing protease inhibitors (Sigma-Aldrich, 5056489001) and benzonase nuclease (Sigma-Aldrich, E1014-25KU). The suspension was centrifuged at 235,000 xg for 0:45 h to collect the insoluble pellet. The pellet was then solubilized in the same buffer containing 2% n-Dodecyl-β-D-Maltopyranoside (DDM) (Anatrace, D310) and stirred for 1 h. The extract was centrifuged at 100,000 xg for 1 h and the supernatant was loaded on Ni-NTA Agarose resin (Qiagen, 30230) pre-equilibrated with 25 mM HEPES, 150 mM NaCl, 50 mM imidazole, pH 7.8, 0.05-0.1% glyco-diosgenin (GDN) (Anatrace, GDN101). The resin was washed with this buffer before elution using buffer containing 500 mM imidazole. The elution fractions were concentrated using 50,000 MWCO concentrators and dialyzed using 20,000 MWCO cassettes (Thermo Fisher Scientific, A52976) against 25 mM HEPES, 150 mM NaCl, pH 8.0, 0.05% GDN. The dialyzed sample was collected and concentrated (50,000 MWCO) further. Protein concentration was estimated by absorbance at 280 nm.

**Purification of Opa1 and Opa1(MGD)**

All steps were done at 4 °C. Harvested *E. coli* cells expressing Opa1(253-960) or Opa1(MGD) were resuspended in 50 mM HEPES, 250 mM NaCl, 1 mM EDTA, pH 8.0, 1 mg/ml lysozyme, protease inhibitors and benzonase nuclease. The cell suspension was lysed by sonication and centrifuged at 30,000 xg for 1 h. The supernatant was flowed over StrepTactin XT 4Flow high-capacity resin (IBA Lifesciences, 2-5030-010) packed in a gravity column and equilibrated with 25 mM HEPES, 150 mM NaCl, pH 8.0. The same buffer was used to wash the resin and the bound protein was eluted using buffer containing 50 mM biotin (IBA Lifesciences, 2-1016-005). The fractions were concentrated (50,000 MWCO for Opa1, 20,000 MWCO for Opa1(MGD)) and dialyzed (20,000 MWCO) against 25 mM HEPES, 150 mM NaCl, pH 8.0. The concentrations were estimated using absorbance at 280 nm. The proteins were aliquoted and frozen in liquid nitrogen and stored at -80 °C.

**Generation of SLC25A46-Opa1 and SLC25A46-Mfn2 complexes**

All purification steps were performed at 4 °C or on ice. To purify SLC25A46-Opa1 complex, *P. pastoris* co-expressing SLC25A46-His and Strep-Opa1 was harvested and cryo-milled as before. The milled cells were resuspended in 25 mM HEPES, 150 mM NaCl, 25 mM imidazole, pH 8.0 containing protease inhibitors and benzonase nuclease. GDN (2%) and 18:1 PA (Avanti Polar Lipids, 840875) (0.1 mg/ml solubilized in GDN) were added and stirred for 2 h. The suspension was centrifuged at 100,000 xg for 1 h. The supernatant was loaded onto a 1 ml HisTrap HP column and washed with 25 mM HEPES, 150 mM NaCl, 25 mM imidazole, pH 8.0, 0.025 mg/ml 18:1 PA and 0.1% GDN and eluted with the same buffer containing 500 mM imidazole. The elution was diluted with StrepTactin binding buffer, 25 mM HEPES, 150 mM NaCl, 1 mM EDTA, pH 8.0, 0.025 mg/ml PA, 0.1% GDN to bring the imidazole concentration below 100 mM and added to 1 ml StrepTactin XT 4Flow resin, washed with the above binding buffer and eluted in buffer containing 50 mM biotin. The elution fractions containing SLC25A46 and Opa1 were combined and concentrated (50,000 MWCO) to ~5.5 mg/ml.

To purify SLC25A46-Mfn2 complex, *P. pastoris* co-expressing SLC25A46-His and Mfn2-Strep was harvested, cryo-milled and resuspended in 25 mM HEPES, 150 mM NaCl, 25 mM imidazole, pH 8.0 with protease inhibitors and benzonase nuclease and centrifuged at 235,000 xg for 45 min to collect the membrane fractions. The membrane fractions were resuspended in the same buffer containing 2% GDN and 0.1 mg/ml PA and stirred for 2 h. The extract was centrifuged and processed using Ni-NTA and StrepTactin columns as described above. The elution fractions from StrepTactin column were concentrated (50,000 MWCO) to ~2.4 mg/ml.

**Binding experiments using purified proteins**

Binding reactions were prepared in 25 mM HEPES, 150 mM NaCl, pH 8.0, 0.1% GDN (binding buffer). Purified Opa1 at 2 µM concentration was incubated with varying concentrations of SLC25A46 at 4 °C for 3 h. The reaction mixtures (100 µl) were added to 50 µl StrepTactin XT 4Flow high-capacity resin packed in spin column and incubated with gentle agitation for 15 min. The spin columns were centrifuged at 500 xg for 30 s to collect the flow-through. The columns were washed thrice with 100 µl of binding buffer. The bound proteins were eluted by incubating with 100 µl of buffer containing 50 mM biotin for 15 min followed by centrifugation at 500 xg for 30 s. Samples analyzed by Western blotting.

**Western blotting**

Following SDS PAGE, proteins were transferred to PVDF membrane (Bio-Rad, 1704156) using Bio-Rad Trans-Blot Turbo Transfer system. The membranes were incubated with the primary antibodies as indicated and were probed with the appropriate secondary antibodies conjugated to horseradish peroxidase. The blots were developed using Western Lightning Plus Chemiluminescent Substrate (Revvity, NEL105001EA) and imaged using Amersham Imager 680. Densitometry was performed using ImageJ. Antibodies: anti-His (Thermo Fisher Scientific, MA1-21315, 1:1000), anti-Strep (IBA Lifesciences, 2-1509-001, ~1:33000), anti-SLC25A46 (Abcam, ab237760, 1:2500), anti-Opa1 (Novus Biologicals, NBP2-59770, 1:1000), anti-Mfn2 (Thermo Fisher Scientific, 702768, 1:1000), anti-MTCO1 (Abcam, ab14705, 1:2000), anti-mouse, HRP conjugated (Sigma-Aldrich, GENA931-1ML, 1:5000), anti-rabbit, HRP conjugated (Sigma-Aldrich, GENA934-1ML, 1:5000).

**Crosslinking mass spectrometry**

For crosslinking, BS3 (Thermo Fisher Scientific, A39266) was added to the concentrated SLC25A46-Opa1(263-960) and SLC25A46-Mfn2 complexes at a final concentration of 0.75 mM. Reactions were incubated on ice for 1 h and quenched by adding Tris-HCl, pH 7.8 at a final concentration of ~45 mM) and incubating 15 min on ice. The samples were separated on a 7.5% polyacrylamide gel and bands excised for mass spectrometry.

While we identified several monolinks and intralinks in Opa1 with BS3 crosslinker, we were unable to identify any crosslinks with SLC25A46 as the sequencing depth for SLC25A46 in this sample was poor. To overcome this, we performed crosslinking experiments with excess purified proteins. We used BS(PEG)_5_ (Thermo Fisher Scientific, 21581) (25 mM stock in 10% DMSO) in addition. The crosslinker (BS3 or BS(PEG)5) was added at a final concentration of 2.5 mM to 20 µM SLC25A46 and 30 µM Opa1(253-960) or Opa1(MGD) and the sample separated on a 12% Bis-Tris 1.0 mm, Mini Protein Gels. The subsequent steps were same as above. The excised gel bands were destained with 1:1 v/v methanol and 100 mM ammonium bicarbonate. Gel pieces were partially dehydrated with acetonitrile then further dehydrated by SpeedVac. Proteins were reduced at 57 ˚C for 1 h by adding 100 µl of 20 mM aqueous DTT solution to the dehydrated gel piece. The solution was removed and replaced with 100 μl of a 50 mM iodoacetamide solution and the gel pieces incubated in the dark for 45 min. the solution was removed, and the gel pieces again partially dehydrated with an acetonitrile rinse and further dried in a SpeedVac. 250 ng of sequencing grade modified trypsin (Promega) was added to each sample followed by 300 µl of 100 mM ammonium bicarbonate to cover the gel pieces. Digestion was allowed to proceeded overnight on a shaker at room temperature. To stop the digestion and assist with peptide extraction a solution of 5% formic acid in acetonitrile 1:2 (v:v) was added and incubated for 15 min with agitation. The solution was transferred into a new tube and the procedure repeated two more times. The combined solutions were dried down in a SpeedVac concentrator to remove the acetonitrile. The samples resuspended in 0.1% acetic acid and loaded onto equilibrated Ultra-Micro SpinColumns™ (Harvard Apparatus) using a microcentrifuge. The spin columns were washed three times with 0.1% trifluoroacetic acid and the last wash with 0.5% acetic acid. Peptides were eluted with 40% acetonitrile in 0.5% acetic acid, followed by 80% acetonitrile in 0.5% acetic acid. The solutions combined and dried down using a SpeedVac concentrator. The samples were reconstituted in 0.5% acetic acid and stored at -80 °C until analysis.

Approximately 1 µg of each sample was analyzed individually by LC/MS/MS. Samples were loaded onto an Acclaim PepMap trap column (2 cm x 75 µm) in line with an EASY-Spray analytical column (50 cm x 75 µm ID PepMap C18, 2 μm bead size) using the autosampler of an EASY-nLC 1200 HPLC (Thermo Scientific). Solvent A was 2% acetonitrile in 0.5% acetic acid and solvent B was 80% acetonitrile in 0.5% acetic acid. The gradient used was the following: 5 min held at 5% solvent B, in 60 min to 35% solvent B, in 10 min to 45% solvent B, and in 10 min to 100% solvent B. Peptides were gradient eluted directly into a Thermo Scientific Orbitrap Eclipse Mass Spectrometer.

Mfn2 and SLC25A46 Crosslink analysis:

High resolution full spectra were acquired with a resolution of 240,000, an AGC target of 1e6, with a maximum ion time of 50 ms, and a scan range of 400 to 1500 m/z. All precursors with charge states between 2-7 were selected for fragmentation with a dynamic exclusion of 20 sec. All MS/MS HCD spectra were collected in the ion trap using the rapid scan mode, an AGC target of 2e4, maximum inject time of 18 ms, one microscan, 0.7 m/z isolation window, and normalized collision energy (NCE) of 27. The instrument was set to run at top speed with a cycle time of 3 sec. Samples were run a second time to specifically only fragment precursor ions that are likely crosslinked (charge state 4-10) using either HCD or ETD fragmentation. The gradient used was the following: 5 min held at 5% solvent B, in 120 min to 35% solvent B, in 10 min to 45% solvent B, and in 10 min to 100% solvent B. High resolution full spectra were acquired with a resolution of 60,000, an AGC target of 4e5, with a maximum ion time of 50 ms, and a scan range of 400 to 1500 m/z. Charge states 4-6 were analyzed in the Orbitrap using HCD with an NCE of 27, resolution of 30K, AGC target of 2e5, maximum inject time of 200 msec and an isolation window of 2. Charge states 7-10 were analyzed in the Orbitrap using ETD with a charge state dependent reaction time, resolution of 30K, AGC target of 2e5, maximum inject time of 54 msec and an isolation window of 2.

Opa1 and SLC25A46 Crosslink analysis:

High resolution full spectra were acquired with a resolution of 120,000, an AGC target of 4e6, with a maximum ion time of 50 ms, and a scan range of 400 to 1500 m/z. All precursors with charge states between 2-5 were selected for fragmentation with a dynamic exclusion of 30 sec. All MS/MS HCD spectra were collected in the Orbitrap with a resolution of 30K, an AGC target of 2e5, maximum inject time of 200 ms, one microscan, 2 m/z isolation window, and normalized collision energy (NCE) of 27.

MS/MS spectra were searched in Byos by Protein Metrics against the human SLC25A46, Opa1 and Mfn2 sequences to identify crosslinked peptides.

**Cell lines and culture conditions**

HCT116 WT and SLC25A46 KO cells were kind gifts from CeMM (Research Center for Molecular Medicine, Medical University of Vienna, Austria). The cells maintained in RPMI 1640 with 10% fetal bovine serum (FBS) and 1% penicillin/streptomycin at 37 °C and 5% CO_2_. Cell genotype confirmed by Western blotting. Human SLC25A46 sequence from plasmid pDONR221_SLC25A46 (a gift from RESOLUTE Consortium & Giulio Superti-Furga (Addgene plasmid #131971; http://n2t.net/addgene:131971; RRID:Addgene_131971) was cloned into pCDNA3.1(+) plasmid to generate the construct pAH9. Site-directed mutagenesis (New England Biolabs) was used to generate expression plasmids bearing the mutations G249D (pAH10), R257Q (pAH11), R340C (pAH12), I349D (pAH13), ∆1-83 (pAH14) and R257A/R347A (pAH36) in the SLC25A46 coding sequence of pAH9. To transiently express SLC25A46 and its mutant derivatives in HCT116 SLC25A46 KO cells, cells seeded onto 35 mm glass-bottom dishes (MatTek Life Sciences) coated with poly-D-lysine (0.1 mg/ml) were transfected with 2.5 µg expression plasmid mixed with Lipofectamine 3000 in a total volume of 250 µl Opti-MEM (Thermo Fisher Scientific). Cells were grown for 36-48 h prior to imaging.

For Western blotting, mitochondria were isolated using Qiagen Qproteome Mitochondria Isolation Kit (Qiagen, 37612). Protease inhibitors were added to isolated mitochondria and solubilized using 0.5% DDM. Protein content was estimated using Bradford assay (Bio-Rad, 5000205). ~1 µg of total protein was loaded on SDS-PAGE gel.

**Fluorescence imaging and quantification**

HCT116 cells were seeded onto glass-bottom dishes and grown to ~80% confluency. Cells were washed with PBS and fixed in 4% paraformaldehyde (PFA) in PBS for 20 min. Following three PBS washes, cells were permeabilized in 0.1% Triton X-100 diluted in PBS for 5 min. Cells were incubated in 2% normal goat serum and 0.05% Triton X-100 in PBS for 1 h, then incubated in the same buffer with 1:300 Tom20 rabbit antibody (Invitrogen, MAS-34964) for 1 h at room temperature or overnight at 4˚C. After 3 washes with PBS, cells were incubated with 1:1000 goat anti-rabbit Alexa Fluor® 488 AffiniPure antibody (Jackson ImmunoResearch, 111-545-144) for 1 h at room temperature. Cells were washed in PBS and coated with UltraCruz® Aqueous Mounting Medium with DAPI (Santa Cruz) prior to imaging. Videos were captured using A1R HD25 point scanning confocal with GaAsP and PMT detectors, equipped with an Apo TIRF 60x/1.49 NA objective lens and Ti2 Z-drive. 300 cells per transfected condition were manually scored into four morphological classifications (1): “Fragmented” refers to cells that contain spherical mitochondrial fragments with two or less short tubules present. “Short tubular” refers to cells containing a mixture of fragmented and short tubular mitochondria. “Long tubular” refers to cells with elongated mitochondria, but not fused into a mitochondrial mesh. “Interconnected” refers to cells with a highly interconnected network of mitochondrial filaments, with few isolated mitochondria.

**Live cell confocal microscopy**

Cells were stained with 1:1000 PKmito Deep Red (Spirochrome) at 37˚C for 30 min. Following three 1X PBS washes, cells were placed in Live Cell Imaging Solution (Invitrogen). Acquisition was performed using Nikon A1R HD25 confocal with GaAsP and PMT detectors and an Apo 60x/1.49 NA objective. Temperature, humidity, and CO_2_ were controlled within an Oko-Lab chamber equipped with Live Cell^TM^ and Solent Scientific controllers. Image acquisition done using NIS-Elements (Nikon Instruments Inc.) and analysis performed using Fiji (2). Time-lapse videos of mitochondria taken at 1 frame per 10 sec for 5 min.

**Supporting Figures**

**Supporting Figure 1. SLC25A46 interacts with Opa1.**

A. Strep-Opa1 co-elutes upon elution of bound SLC25A46-His. GDN extracts prepared from *P. pastoris* expressing SLC25A46-His, Strep-Opa1 or co-expressing SLC25A46-His and Strep-Opa1 were subjected to Ni-NTA column binding and elution.

B. The intralinks (black arcs) and monolinks (red circles) identified from crosslinking-mass spectrometry experiments are mapped on the domain diagrams of SLC25A46 and Opa1.

C. AlphaFold2 model for SLC25A46-Opa1 complex colored by pLDDT (red = high confidence, blue = low confidence).

**Supporting Figure 2. SLC25A46 mutations in the binding interface diminish binding to Opa1.**

A. Intensity of the anti-His (SLC25A46) bands in the elution in Figure 2 were normalized against the corresponding bands from the input and plotted as a function of SLC25A46 concentration used in the reaction mixtures.

B. Presence of the nucleotides did not affect binding of SLC25A46 to Opa1. GDN extracts prepared from *P. pastoris* co-expressing SLC25A46-His and Strep-Opa1 were incubated with 2 mM of the indicated guanosine nucleotide prior to StrepTactin binding and elution.

**Supporting Figure 3. The BSE of Opa1 is sufficient to bind SLC25A46.**

A. Purified SLC25A46-His WT was incubated with 2 µM of Strep-Opa1 or Strep-Opa1(MGD) and subjected to StrepTactin binding and elution. Input and elution analyzed by Western blotting. Control reaction: 10 µM of SLC25A46-His alone.

B. Circular map of SLC25A46 and Opa1(MGD) with identified lysine-lysine crosslinks (black arcs) shown. Crosslinker used is indicated. Crosslinks were identified in BSE helices 2 and 3 of Opa1(MGD) as in Opa1.

C. A model of the SLC25A46-Opa1(MGD) interaction. The N-terminal region of SLC25A46 (residues 1-83) is disordered and not displayed. Boxed region enlarged on right displaying the identified crosslinks and the calculated Cɑ-Cɑ distances based on the model.

**Supporting Figure 4. SLC25A46 interacts with Mfn2.**

A. Mfn2-Strep co-elutes upon elution of bound SLC25A46-His. GDN extracts prepared from *P. pastoris* expressing Mfn2-Strep or co-expressing SLC25A46-His and Mfn2-Strep were subjected to Ni-NTA column binding and elution.

B. GDN extracts prepared from *P. pastoris* co-expressing SLC25A46-His and Mfn2-Strep were incubated with 2 mM of the indicated guanosine nucleotide prior to StrepTactin binding and elution. Presence of the nucleotides did not affect binding of SLC25A46 to Mfn2.

C. The intralinks (black arcs) and monolinks (red circles) identified from crosslinking-mass spectrometry experiments are mapped on the domain diagram of Mfn2.

D. Alphafold2 model of Mfn2 colored by pLDDT (red = high confidence, blue = low confidence).

**Supporting Figure 5.**

A. GDN extracts from *P. pastoris* co-expressing Strep-Opa1 and SLC25A46-His WT or mutants subjected to StrepTactin (middle) or Ni-NTA (bottom) column binding and elution.

B. GDN extracts prepared from *P. pastoris* co-expressing Mfn2-Strep and either SLC25A46-His WT, (Δ2-83)SLC25A46-His or disease mutants were subjected to StrepTactin (middle) or Ni-NTA (bottom) column binding and elution. (Δ2-83)SLC25A46 still interacts with Mfn2 as seen by co-elution of both proteins. Disease mutations did not abrogate binding of SLC25A46 to Opa1 or Mfn2.

**Supporting Video 1.**

Live-cell confocal microscopy of PKmito Deep Red-stained mitochondria in indicated HCT116 cell lines. Movies were taken at 15 sec per frame for 5 min. Playback at 4 frames per sec (37.5x real-time). Scale bar = 10 µm.

**Supporting Figure 1**

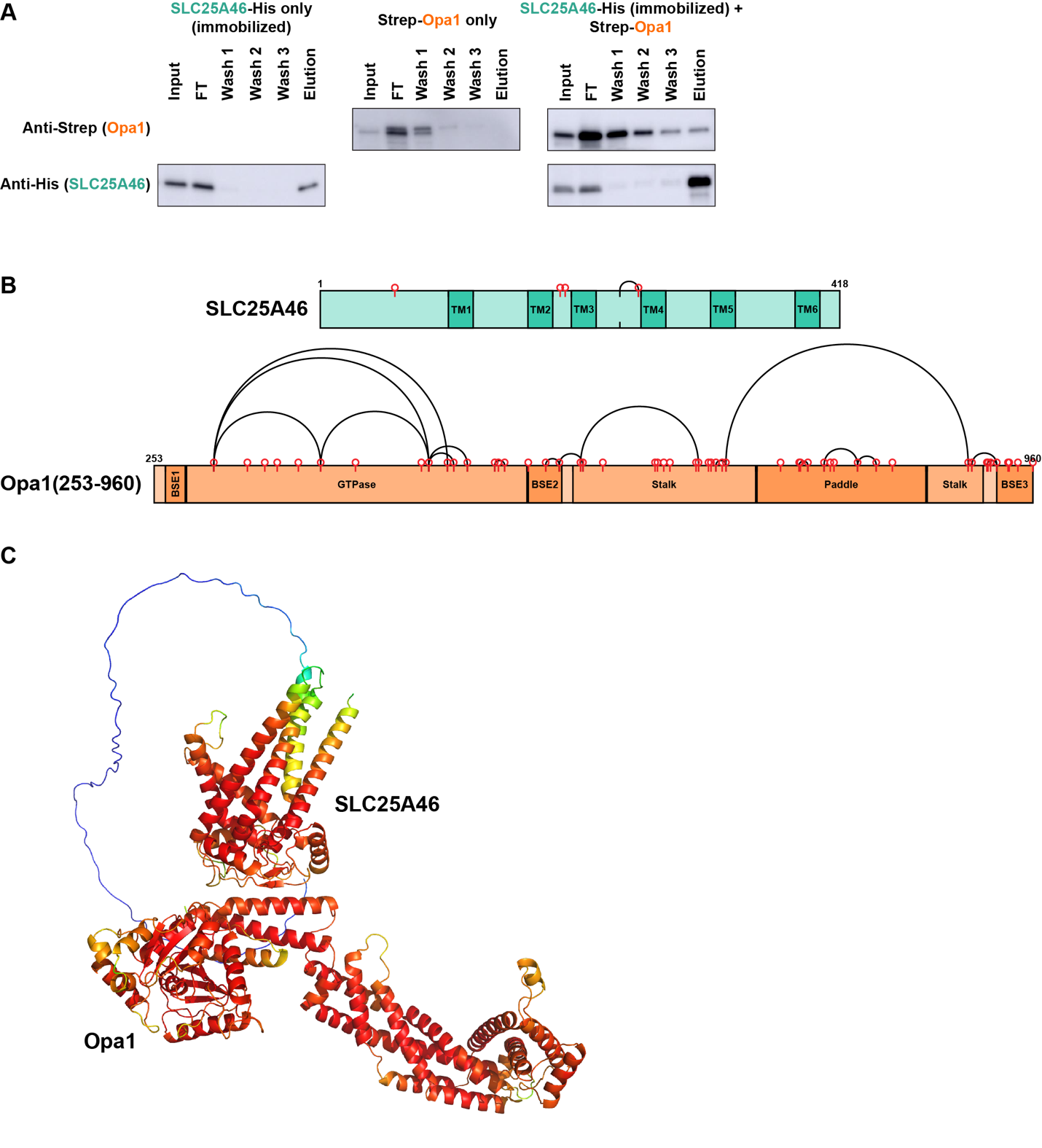

**Supporting Figure 2**

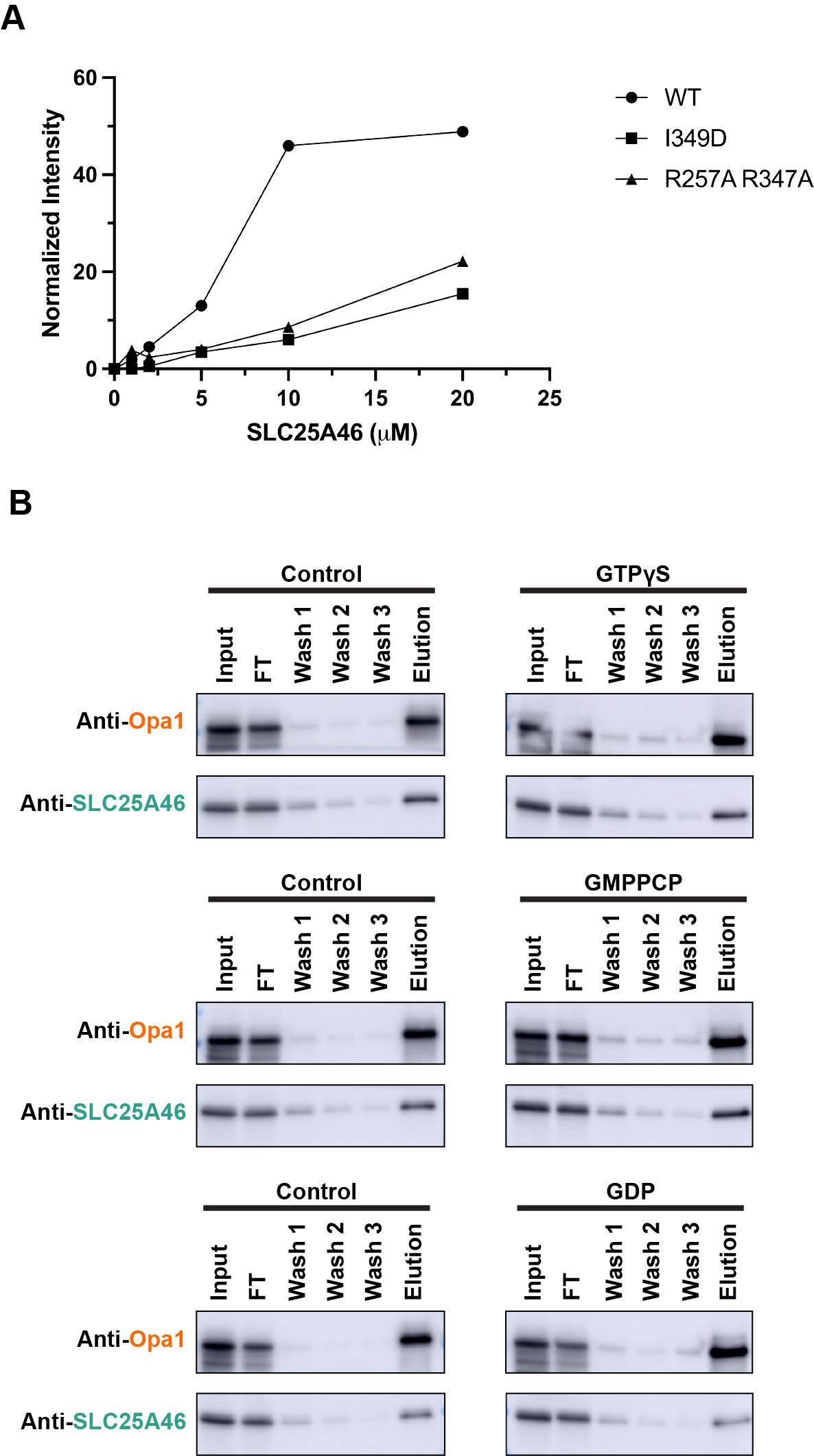

**Supporting Figure 3**

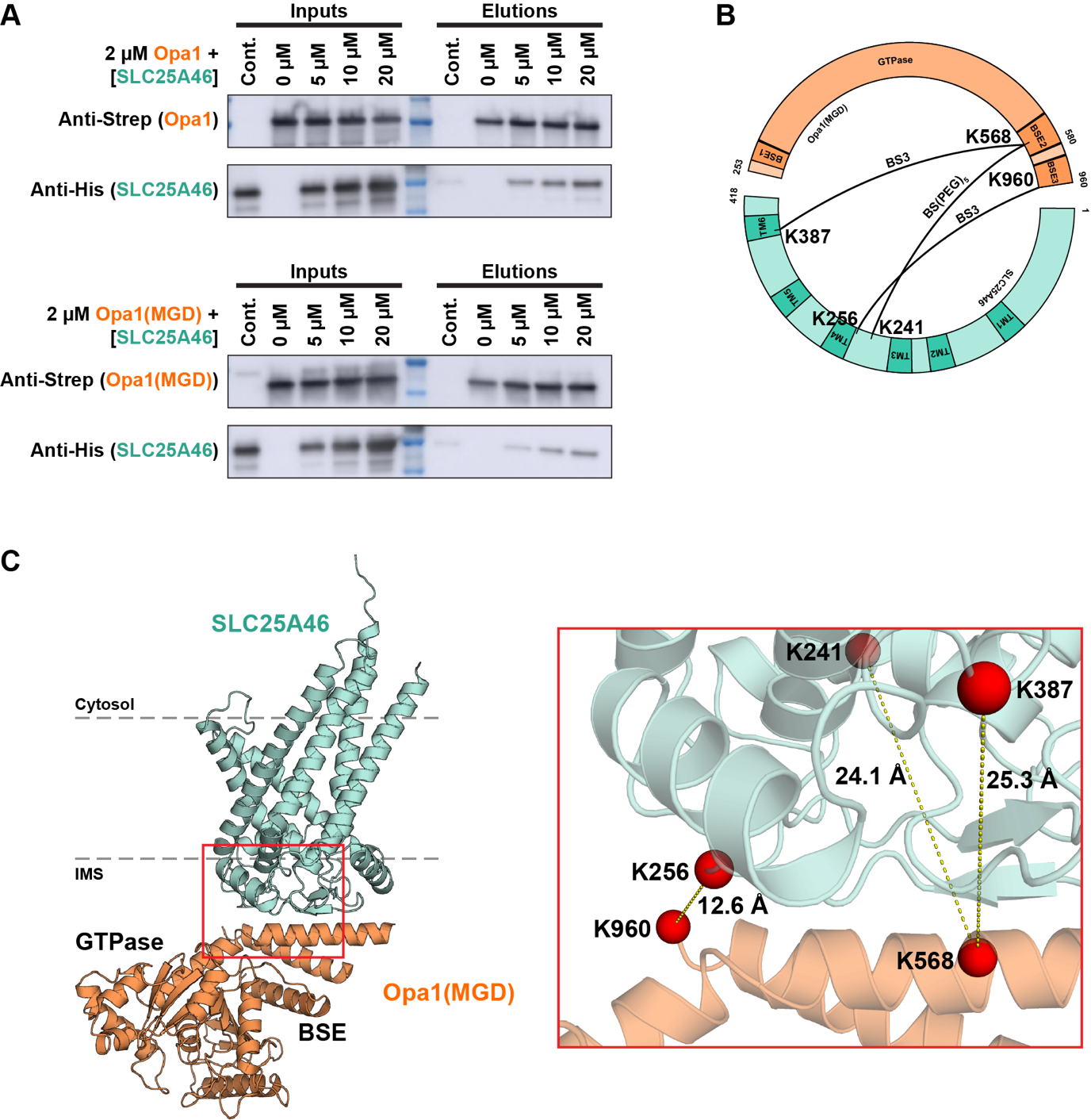

**Supporting Figure 4**

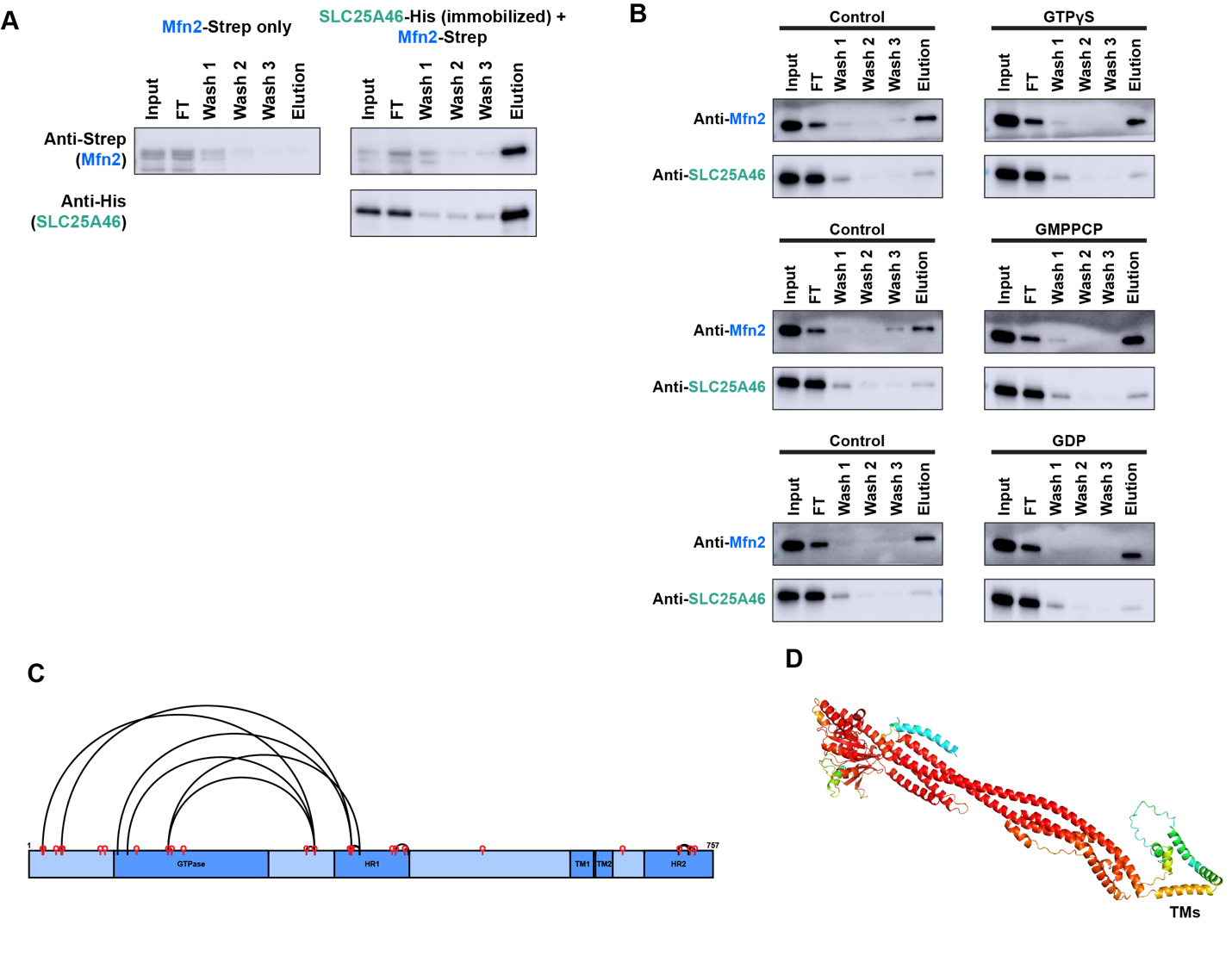

**Supporting Figure 5**

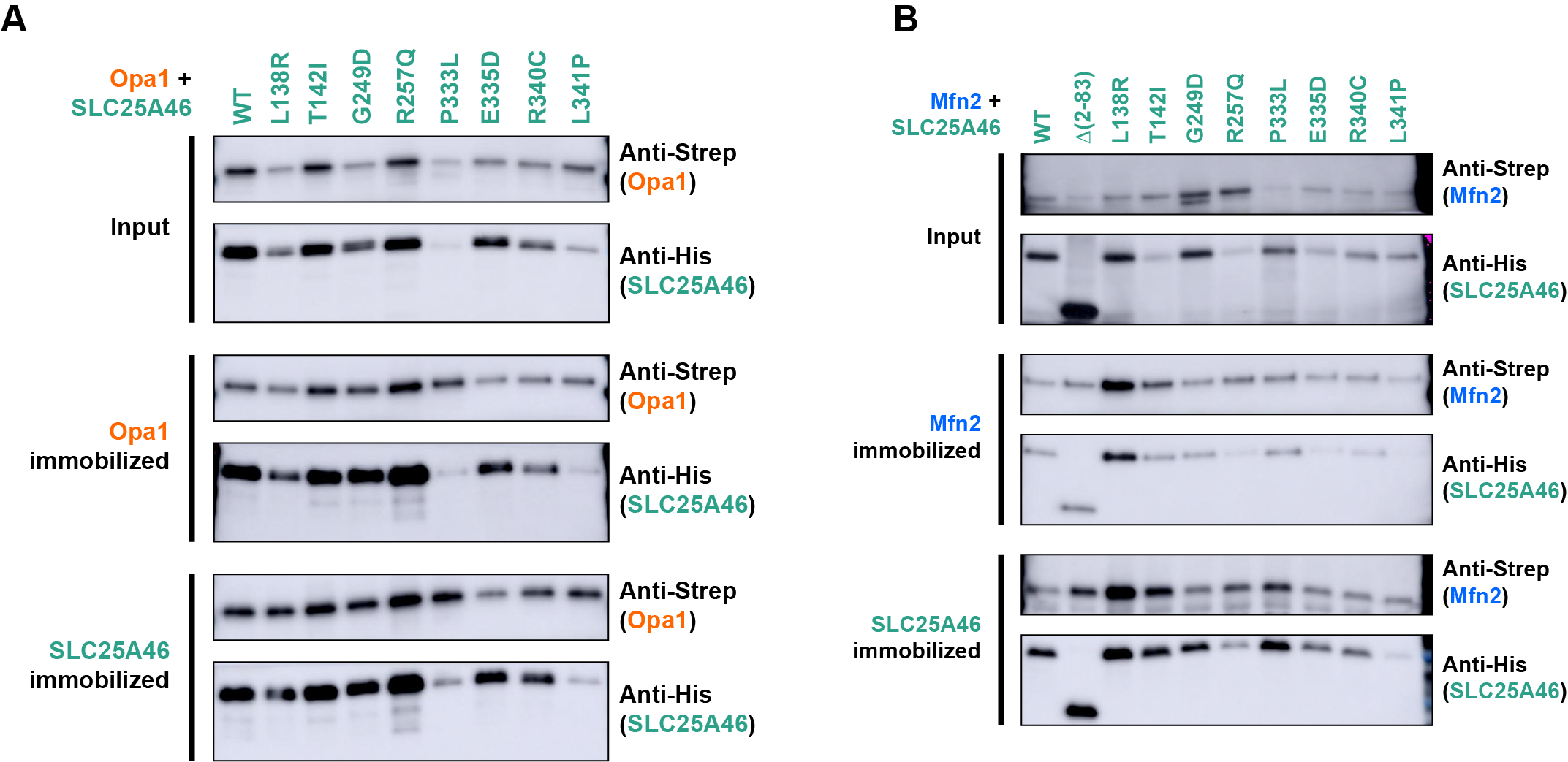
